# Spatiotemporal context modulates encoding and retrieval of overlapping events

**DOI:** 10.1101/2021.05.17.444541

**Authors:** Devyn E. Smith, Isabelle L. Moore, Nicole M. Long

## Abstract

Overlap between events can lead to interference due to a tradeoff between encoding the present event and retrieving the past event. Temporal context information – ‘when’ something occurred, a defining feature of episodic memory – can cue retrieval of a past event. However, the influence of temporal overlap, or proximity in time, on the mechanisms of interference are unclear. Here, by identifying brain states using scalp electroencephalography (EEG) from male and female human subjects, we show the extent to which temporal overlap promotes interference and induces retrieval. In this experiment, subjects were explicitly directed to either encode the present event or retrieve a past, overlapping event while perceptual input was held constant. We find that the degree of temporal overlap between events leads to selective interference. Specifically, greater temporal overlap between two events leads to impaired memory for the past event selectively when the top-down goal is to encode the present event. Using pattern classification analyses to measure neural evidence for a retrieval state, we find that greater temporal overlap leads to automatic retrieval of a past event, independent of top-down goals. Critically, the retrieval evidence we observe likely reflects a general retrieval mode, rather than retrieval success or effort. Collectively, our findings provide insight into the role of temporal overlap on interference and memory formation.

## Introduction

Overlap between events leads to interference and impairs memory for those events (McGeoch, 1942; Anderson, 2003). For example, at a conference you may talk to a colleague whom you had previously met over Zoom. Later you may have difficulty remembering either the original Zoom meeting or the subsequent conference conversation. The overlap between these events (e.g. the colleague) promotes retrieval of the past event (the meeting on Zoom) while you are trying to encode the present event (your conversation; Kuhl, Shah, DuBrow, & Wagner, 2010). As retrieval and encoding recruit distinct neural substrates and cannot be engaged in simultaneously (Hasselmo, Bodelon, & Wyble, 2002), retrieving the past comes at the expense of encoding the present (Long & Kuhl, 2019). Although overlap is a critical factor in retrieval-mediated interference, two events may overlap along many dimensions and to varying degrees. Temporal overlap, or proximity in time, has been shown to enhance inference (Zeithamova & Preston, 2017), but it is unclear how temporal overlap contributes to interference. The aim of this study is to investigate the extent to which temporal overlap induces retrieval and, in turn, impacts interference.

Temporal information is a hallmark of episodic memory (Tulving, 1993) and is well known to impact how events are encoded and retrieved. The closer two events are in time and/or space the more likely they are to be recalled together (Kahana, 1996; Manning, Polyn, Baltuch, Litt, & Kahana, 2011) and the greater their neural similarity (Manns, Howard, & Eichenbaum, 2007; Folkerts, Rutishauser, & Howard, 2018). Retrieved context theory (Howard & Kahana, 2002; Sederberg, Howard, & Kahana, 2008; Polyn, Norman, & Kahana, 2009; Lohnas & Kahana, 2014) provides an account for these effects whereby spatiotemporal context – an amalgamation of external stimuli and internal states – is bound, via the hippocampus, to the present experience (Eichenbaum, 2004; Wang & Diana, 2017; Long & Kahana, 2019; Yonelinas, Ranganath, Ekstrom, & Wiltgen, 2019) and is later used by the hippocampus as a cue to retrieve past experiences (Long et al., 2017). Comparison of activity patterns between study and test items – a recalled word or recognition probe – provides support for retrieved context theory in that the shorter the temporal distance between two items at study, the greater the pattern similarity between the study pattern of one item and the test pattern of the other item (Manning et al., 2011; Howard, Viskontas, Shankar, & Fried, 2012; El-Kalliny et al., 2019). Although contextually-mediated retrieval is typically considered in relation to the test phase of an experiment, in principle contextually-mediated retrieval should occur whenever there is a spatiotemporal contextual overlap between items. Such retrieval may occur automatically, or independent from top-down demands (Smith, Handy, Hernandez, & Jacoby, 2018). Therefore, we hypothesized that overlap in spatiotemporal context between two events produces retrieval during study and in turn promotes interference.

Here, we report a human scalp electroencephalography (EEG) study in which subjects studied two sets of object images in which the second set categorically overlapped with the first set. During study of the second set of object images, subjects were explicitly instructed to either encode the second (present) object or retrieve the first (past) object. These instructions were intended to bias subjects toward either an encoding or retrieval state. Our critical manipulation was the temporal distance between the first and second object, whereby the shorter the temporal distance between two objects, the greater their spatiotemporal contextual overlap. Following study, subjects completed a recognition task to probe their memory for all previously-presented objects. To the extent that spatiotemporal contextual overlap influences interference, we should find that temporal distance modulates memory performance for the first and/or second objects. To the extent that spatiotemporal contextual overlap promotes retrieval, we should find that subjects are biased toward a retrieval state during second objects that are presented near in time to a categorically overlapping first object.

## Materials and Methods

### Subjects

Forty (34 female; age range = 18-37, mean age = 20.3 years) right-handed, native English speakers from the University of Virginia community participated. All subjects had normal or corrected-to-normal vision. Informed consent was obtained in accordance with the University of Virginia Institutional Review Board for Social and Behavioral Research and subjects were compensated for their participation. Three subjects were excluded from the final dataset: one who previously completed a behavioral version of the task, one who had poor task performance (recognition accuracy below three standard deviations of the mean of the full dataset), and one due to technical issues resulting in poor signal quality throughout the majority of the session. Thus, data are reported for the remaining 37 subjects. The raw, de-identified data and the associated experimental and analysis codes used in this study will be made available via the Long Term Memory laboratory website upon publication.

### Mnemonic State Task Experimental Design

Stimuli consisted of 576 object pictures, drawn from an image database with multiple exemplars per object category (Konkle, Brady, Alvarez, & Oliva, 2010). From this database, we chose 144 unique object categories and 4 exemplars from each category. For each subject, one exemplar in a set of four served as a List 1 object, one as a List 2 object, and the two remaining exemplars served as lures for the recognition phase. Object condition assignment was randomly generated for each subject.

#### General Overview

In each of eight runs, subjects viewed two lists containing 18 object images. For the first list, each object was new (List 1 objects). For the second list (List 2 objects), each object was again new, but was categorically related to an object from the first list. For example, if List 1 contained an image of a bench, List 2 would contain an image of a different bench (Figure 1). During List 1, subjects were instructed to encode each new object. During List 2, however, each trial contained an instruction to either encode the current object (e.g., the new bench) or to retrieve the corresponding object from List 1 (the old bench). The critical manipulation was the distance between the corresponding List 1 and List 2 objects. We divided each list of 18 objects into thirds according to serial position (first [1-6], middle [7-12], and last [13-18]). The objects in the first third of List 1 were “paired” with the objects in the last third of List 2. For example, if List 1 contained an image of a bench in serial position 1, List 2 would contain an image of a different bench in serial position 13-18. The objects in the middle third of List 1 were paired with the objects in the middle third of List 2. The objects in the last third of List 1 were paired with the objects in the first third of List 2. We coded List 1 and List 2 objects as *near* and *far* based on the lag, or difference in serial position, between the two objects in a pair. List 1 and List 2 objects separated by fewer than 18 intervening objects were coded as *near*; List 1 and List 2 objects separated by 18 or more intervening objects were coded as *far*. Following eight runs, subjects completed a two-alternative forced-choice recognition test that separately assessed memory for List 1 and List 2 objects.

**Figure 1.**
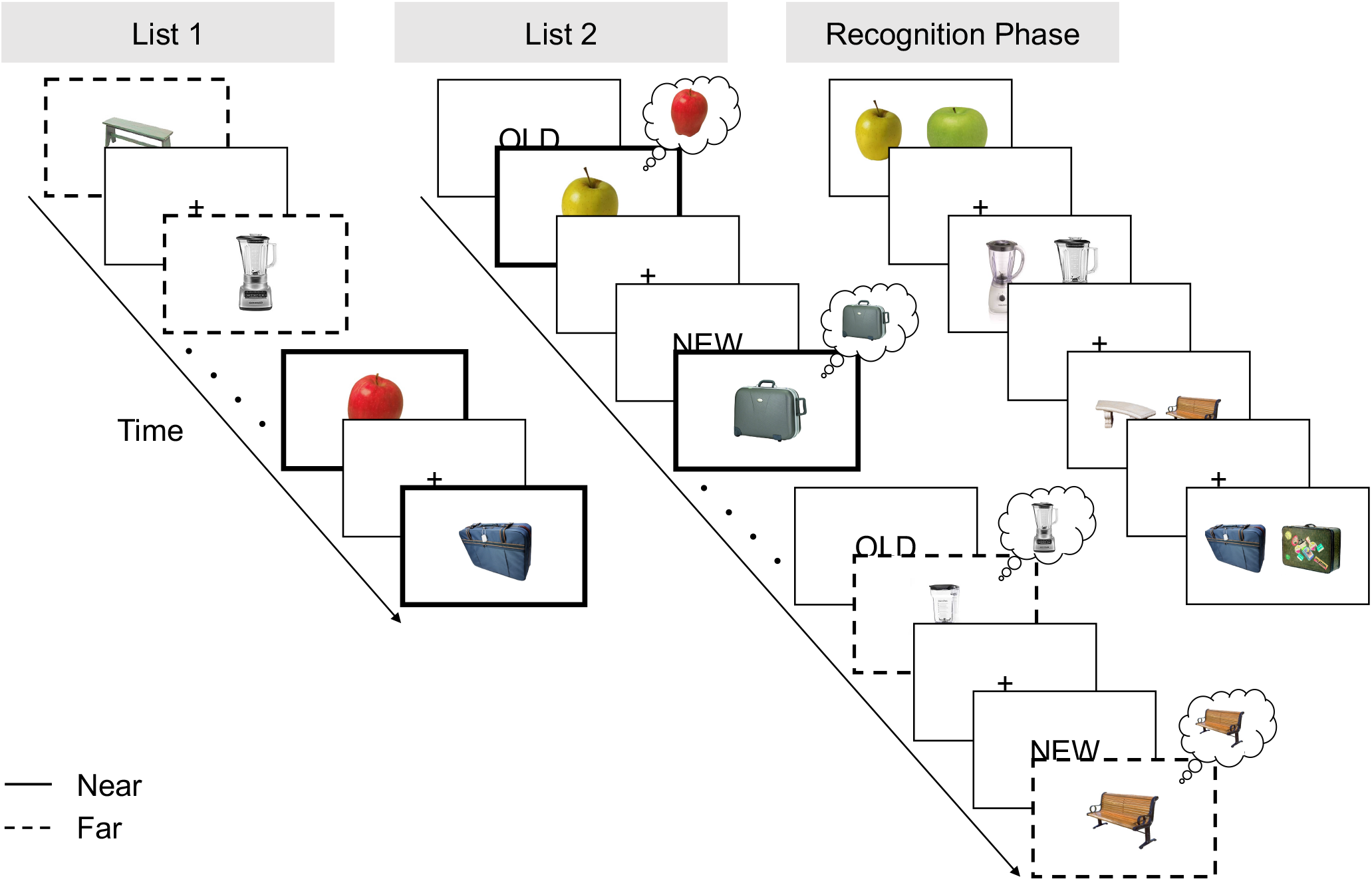
Task Design. During List 1, subjects studied individual objects (e.g. bench, apple). During List 2, subjects saw novel objects that were from the same categories as the objects shown in List 1 (e.g., a new bench, a new apple). Preceding each List 2 object was an “OLD” instruction cue or “NEW” instruction cue. The “OLD” cue signaled that subjects were to *retrieve* the corresponding object from List 1 (e.g., the old apple). The “NEW” cue signaled that subjects were to *encode* the current object (e.g. the new bench). Each run of the experiment contained a List 1 and List 2; object categories (e.g., bench) were not repeated across runs. List 1 and List 2 objects separated by fewer than 18 intervening objects were coded as *near* and List 1 and List 2 objects separated by 18 or more intervening objects were coded as *far* (see Methods). Lines around the boxes are shown for illustrative purposes and were not present during the actual experiment. After eight runs, subjects completed a two alternative forced choice recognition test that tested memory for each List 1 and List 2 object. On each trial, a previously presented object, either from List 1 or List 2, was shown alongside a novel lure from the same category. The subject’s task was to choose the previously presented object. List 1 and List 2 objects were never presented together.

#### List 1

On each trial, subjects saw a single object presented for 2000 ms followed by a 1000 ms inter-stimulus interval (ISI). Subjects were instructed to study the presented object in anticipation for a later memory test.

#### List 2

On each trial, subjects saw a cue word, either “OLD” or “NEW” for 2000 ms. The cue was followed by presentation of an object for 2000 ms, which was followed by a 1000 ms ISI. All objects in List 2 were non-identical exemplars drawn from the same category as the objects presented in the immediately preceding List 1. That is, if a subject saw a bench and an apple during List 1, a different bench and a different apple would be presented during List 2. On trials with a “NEW” instruction (encode trials), subjects were to encode the presented object. On trials with an “OLD” instruction (retrieve trials), subjects tried to retrieve the categorically related object from the preceding List 1. Importantly, this design prevented subjects from completely ignoring List 2 objects following “OLD” instructions in that they could only identify the to-be-retrieved object category by processing the List 2 object.

Subjects completed eight runs with two lists in each run (List 1, List 2). Subjects viewed 18 objects per list, yielding a total of 288 object stimuli from 144 unique object categories. Subjects did not make a behavioral response during either List 1 or 2. Following the eight runs, subjects completed a two-alternative forced choice recognition test.

#### Recognition Phase

Following the eight runs, subjects completed the recognition phase. On each trial, subjects saw two exemplars from the same object category (e.g. two benches; Figure 1). One object had previously been encountered either during List 1 or 2. The other object was a lure and had not been presented during the experiment. Trials were self-paced and subjects selected (via button press) the previously presented object. Trials were separated by a 1000 ms ISI. There were a total of 288 recognition trials (corresponding to the 288 total List 1 and 2 objects presented in the experiment). Note: List 1 and List 2 objects never appeared in the same trial together, thus subjects never had to choose between two previously presented objects. List 1 and List 2 objects were presented randomly throughout the test phase.

### EEG Data Acquisition and Preprocessing

EEG recordings were collected using a BrainVision system and an ActiCap equipped with 64 Ag/AgCl active electrodes positioned according to the extended 10-20 system. All electrodes were digitized at a sampling rate of 1000 Hz and were referenced to electrode FCz. Offline, electrodes were later converted to an average reference. Impedances of all electrodes were kept below 50 kΩ. Electrodes that demonstrated high impedance or poor contact with the scalp were excluded from the average reference. Bad electrodes were determined by voltage thresholding (see below).

Custom Python codes were used to process the EEG data. We applied a high pass filter at 0.1 Hz, followed by a notch filter at 60 Hz and harmonics of 60 Hz to each subject’s raw EEG data. We then performed three preprocessing steps (Nolan, Whelan, & Reilly, 2010) to identify electrodes with severe artifacts. First, we calculated the mean correlation between each electrode and all other electrodes as electrodes should be moderately correlated with other electrodes due to volume conduction. We *z*-scored these means across electrodes and rejected electrodes with *z*-scores less than-3. Second, we calculated the variance for each electrode as electrodes with very high or low variance across a session are likely dominated by noise or have poor contact with the scalp. We then *z*-scored variance across electrodes and rejected electrodes with a |*z*| *>* = 3. Finally, we expect many electrical signals to be autocorrelated, but signals generated by the brain versus noise are likely to have different forms of autocorrelation. Therefore, we calculated the Hurst exponent, a measure of long-range autocorrelation, for each electrode and rejected electrodes with a |*z*| *>* = 3. Electrodes marked as bad by this procedure were excluded from the average re-reference. We then calculated the average voltage across all remaining electrodes at each time sample and re-referenced the data by subtracting the average voltage from the filtered EEG data. We used wavelet-enhanced independent component analysis (Castellanos & Makarov, 2006) to remove artifacts from eyeblinks and saccades.

### EEG Data Analysis

We applied the Morlet wavelet transform (wave number 6) to the entire EEG time series across electrodes, for each of 46 logarithmically spaced frequencies (2-100 Hz; Long & Kahana, 2015). After log-transforming the power, we downsampled the data by taking a moving average across 100 ms time intervals from 4000 ms preceding to 4000 ms following object presentation, and sliding the window every 25 ms, resulting in 317 time intervals (80 non-overlapping). Power values were then *z*-transformed by subtracting the mean and dividing by the standard deviation power. Mean and standard deviation power were calculated across all first and second objects and across time points for each frequency.

### Univariate Analyses

To test the effect of serial position, our two conditions of interest were primacy objects (objects in serial positions 1-9), and recency objects (objects in serial positions 10-18). Our two contrasts were between primacy and recency List 1 objects and primacy and recency List 2 objects. For each contrast, subject, electrode and frequency, we calculated *z*-transformed power in each of two conditions, averaged over the 2000 ms stimulus interval.

### Pattern Classification Analyses

Pattern classification analyses were performed using penalized (L2) logistic regression (penalty parameter = 1), implemented via the sklearn linear model module in Python. Before pattern classification analyses were performed, an additional round of *z*-scoring was performed across features to eliminate trial-level differences in spectral power (Kuhl & Chun, 2014; Long & Kuhl, 2018). Therefore, mean univariate activity was matched precisely across all conditions and trial types. Classifier performance was assessed in two ways. “Classification accuracy” represented a binary coding of whether the classifier successfully guessed the instruction condition. We used classification accuracy for general assessment of classifier performance (i.e., whether encode/retrieve instructions could be decoded). “Classifier evidence” was a continuous value reflecting the logit-transformed probability that the classifier assigned the correct instruction for each trial. Classifier evidence was used as a trial-specific, continuous measure of mnemonic state information, which was used to assess the degree of retrieval evidence present on *near* and *far* trials.

We trained within-subject classifiers to discriminate List 2 encode vs. retrieve trials based on a feature space comprised of all 63 electrodes *×* 46 logarithmically spaced frequencies ranging from 2 to 100 Hz. For each subject, we used leave-one-run-out cross validated classification in which the classifier was trained to discriminate encode from retrieve instructions for seven of the eight runs and tested on the held-out run. For classification analyses in which we assessed classifier accuracy, we averaged spectral power over the 2000 ms stimulus interval. For analyses measuring classifier evidence, we averaged spectral power over four separate 500-ms time intervals across the 2000 ms stimulus interval.

To measure the ability of the classifier to generalize across temporal distance, we trained and tested two separate classifiers to distinguish List 2 encode/retrieve trials. One classifier was trained on *near* trials and tested on *far* trials, the other classifier was trained on *far* trials and tested on *near* trials. As there was a slight imbalance in the number of encode and retrieve trials within each distance, we subsampled trials from the condition with the greater number of trials to match the condition with fewer trials. We repeated this procedure for 100 iterations and averaged the resulting classification accuracy values across the 100 iterations.

### Statistical Analyses

We used repeated measures ANOVAs and paired-sample *t*-tests to assess the effect of instruction (encode, retrieve) and temporal distance (*near, far*) on behavioral memory performance.

We used paired-sample *t*-tests to compare classification accuracy across subjects to chance decoding accuracy, as determined by permutation procedures. Namely, for each subject we shuffled the condition labels of interest (e.g., encode and retrieve for the List 2 instruction classifier) and then calculated classification accuracy. We repeated this procedure 1000 times for each subject and then averaged the 1000 shuffled accuracy values for each subject. These mean values were used as subject-specific empirically derived measures of chance accuracy.

We used repeated measures ANOVAs and paired-sample *t*-tests to assess the interaction between instruction (encode, retrieve), temporal distance (*near, far*), and time interval on retrieval evidence.

## Results

### Influence of Spatiotemporal Contextual Overlap on Interference

We first sought to replicate the finding that subjects are able to shift between encoding and retrieval states in a goal directed manner (Long & Kuhl, 2019), by testing whether instructions influenced performance on the recognition task. Although encode/retrieve instructions only appeared during List 2, we also considered whether memory for List 1 objects was influenced by List 2 instructions (e.g., whether memory for the old bench was influenced by whether the new bench was associated with an encode vs. retrieve instruction). A two-way, repeated measures ANOVA with factors of list (1, 2) and instruction (encode, retrieve) revealed a list by instruction interaction (*F*_1,36_ = 6.045, *p* = 0.0189; Figure 2A). This interaction was driven by greater recognition for List 2 objects presented with an encode (M = 82.88%, SD = 8.51%) relative to a retrieve instruction (M = 80.52%, SD = 7.79%; difference between List 2 encode vs retrieve: *t*_36_ = 2.1072, *p* = 0.0421) and numerically greater recognition for List 1 objects presented with a retrieve (M = 84.27%, SD = 7.7%) relative to an encode instruction (M = 83.3%, SD = 7.03%; difference between List 1 encode vs retrieve: *t*_36_ = -1.7542, *p* = 0.0879).

**Figure 2.**
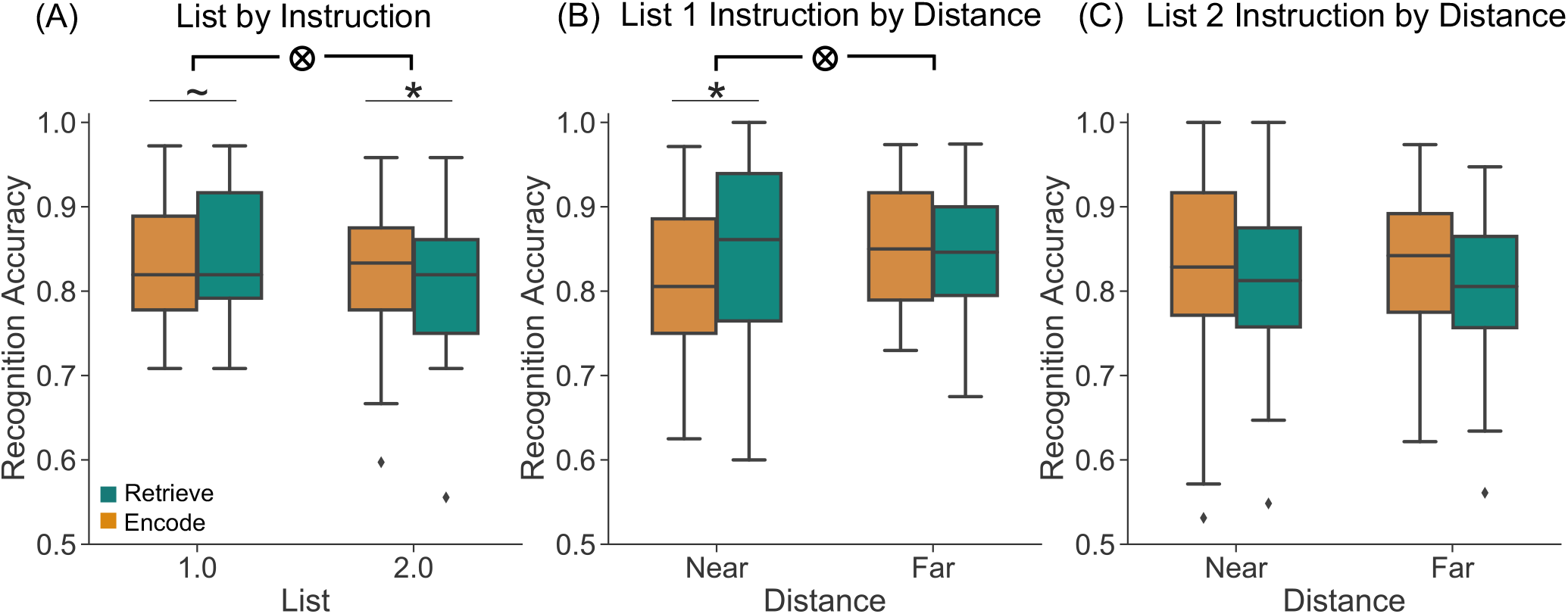
Behavioral Results. We assessed recognition accuracy as a function of list (1, 2), instruction (encode, orange; retrieve, teal), and distance (*near, far*). **(A)** When recognition accuracy is collapsed across distance, we find a significant interaction between list and instruction (*p* = 0.0189) driven by greater accuracy for List 2 objects presented with an encode compared to a retrieve instruction and numerically greater accuracy for List 1 objects presented with a retrieve compared to an encode instruction. **(B)** For recognition accuracy of List 1 objects, we find a significant interaction between instruction and distance (*p* = 0.0435) driven by greater accuracy for *near* retrieve trials compared to *near* encode trials. **(C)** For recognition accuracy of List 2 objects, we find a trending main effect of instruction (*p* = 0.0569) driven by greater accuracy for encode compared to retrieve trials. *∼* p *<* 0.1, * p *<* 0.05.

Having replicated our previous finding that instructions to encode and retrieve modulate behavior, we next sought to test the effect of temporal distance on recognition accuracy, separately for each list. If *near* List 1 objects are more readily retrievable due to temporal overlap with their associated List 2 objects, this may yield different memory outcomes for List 1 and List 2 objects. First, a shorter temporal distance may facilitate List 1 memory specifically for retrieve trials. That is, if *near* List 1 objects are more retrievable, the retrieve instruction during List 2 may be more effective for *near* than *far* objects, providing the subject another opportunity to process the *near* List 1 object. Second, a shorter temporal distance may impair List 1 memory specifically for encode trials. The intuition is that automatically retrieved *near* List 1 objects may be inhibited or suppressed by virtue of being goal-irrelevant during encode trials. This outcome would be analogous to the inhibition that is thought to occur during retrieval induced forgetting (Anderson, Bjork, & Bjork, 1994; Anderson, 2003). Finally, a shorter temporal distance may impair List 2 memory regardless of instruction, as retrieval driven by the *near* List 1 objects will tradeoff with encoding of the List 2 objects.

We assessed whether the distance between objects, as well as the instruction given during List 2, influenced recognition memory separately for List 1 objects (Figure 2B) and List 2 objects (Figure 2C). For List 1, a two-way, repeated measures ANOVA with factors of instruction (encode, retrieve) and distance (*near, far*), revealed a significant main effect of distance (*F*_1,36_ = 4.931, *p* = 0.0330) driven by greater recognition accuracy for *far* compared to *near* objects. There was a trending main effect of instruction (*F*_1,36_ = 3.769, *p* = 0.0601) driven by greater recognition accuracy for retrieve compared to encode instructions. There was a significant interaction between instruction and distance (*F*_1,36_ = 4.381, *p* = 0.0435), driven by greater accuracy for *near* retrieve trials (M = 83.88%, SD = 11.11%) relative to *near* encode trials (M = 80.9%, SD = 9.17%; difference between *near* encode vs *near* retrieve: *t*_36_ = -2.6225, *p* = 0.0127). Notably, recognition accuracy on *near* encode trials was significantly worse compared to both *far* encode trials (*t*_36_ = -3.3417, *p* = 0.0020) and *far* retrieve trials (*t*_36_ = -3.1204, *p* = 0.0035). For List 2, a two-way, repeated measures ANOVA with factors of instruction (encode, retrieve) and distance (*near, far*), revealed no main effect of distance (*F*_1,36_ = 0.0, *p* = 0.99), and a trending main effect of instruction (*F*_1,36_ = 3.87, *p* = 0.0569) driven by greater recognition accuracy for encode compared to retrieve instructions. The interaction between instruction and distance was not significant (*F*_1,36_ = 0.296, *p* = 0.5900).

We observed decreased recognition accuracy for List 1 *near* objects when subjects attempted to encode the List 2 object compared to when they attempted to retrieve the *near* List 1 object. In fact, *near* List 1 objects paired with the encode instruction are remembered worse than all other List 1 objects, strongly suggesting that bottom-up or automatic retrieval of the *near* List 1 object, when coupled with the top-down demand to encode the List 2 object, leads to suppression of the *near* List 1 object. In comparison, recognition accuracy for List 2 objects was influenced by instruction but not distance, suggesting that subjects were able to flexibly shift between encoding and retrieval states regardless of temporal overlap.

### Influence of Spatiotemporal Contextual Overlap on Retrieval State

Our first goal was to replicate our previous finding that a pattern classifier trained on spectral signals can distinguish encode and retrieve trials (Long & Kuhl, 2019). We conducted a multivariate pattern classification analysis in which we trained a classifier to discriminate encode vs. retrieve List 2 trials based on a feature space comprised of all 63 electrodes and 46 frequencies ranging from 2 to 100 Hz. For this analysis, we averaged spectral power over the 2000 ms stimulus interval. Using within-subject, leave-one-run-out classifiers, mean classification accuracy was 53.55% (SD = 7.81%), which was significantly greater than chance, as determined by permutation tests (*t*_36_ = 2.7241, *p* = 0.0099; Figure 3A).

**Figure 3.**
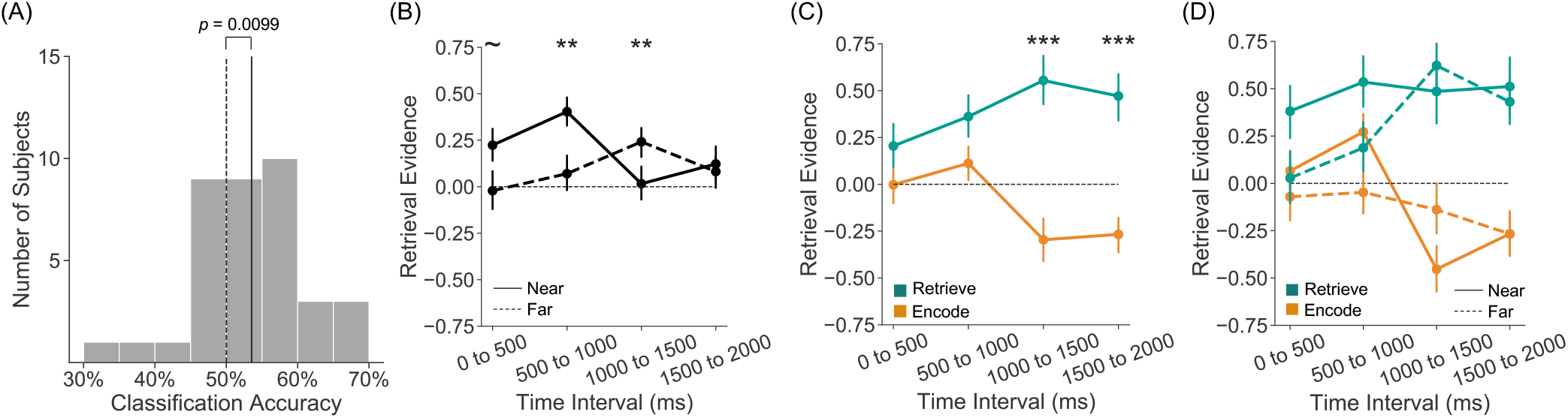
Retrieval State Evidence. We trained an L2-logistic regression classifier to discriminate encode vs. retrieve trials during List 2. The classifier was trained and tested on spectral power across 63 electrodes and 46 frequencies. **(A)** The classifier was trained on average spectral power across the 2000 ms stimulus interval. Mean classification accuracy across all subjects (solid vertical line) is shown along with a histogram of classification accuracies for individual subjects (gray bars) and mean classification accuracy for permuted data across all subjects (dashed vertical line). Mean classification accuracy for permuted data ranged from 49.75% to 50.22% across individual subjects (1000 permutations per subject). Mean classification accuracy was 53.55%, which differed significantly from chance (*p* = 0.0099). **(B-D)** We trained and tested four classifiers on four 500 ms time intervals within the 2000 ms stimulus interval. **(B)** When we average retrieval evidence over instruction, we find a significant interaction between distance and time interval (*p* = 0.0013) driven by greater retrieval evidence on *near* compared to *far* trials early in the stimulus interval. **(C)** When we average retrieval evidence over distance, we find a significant interaction between instruction and time interval (*p* = 0.0024) driven by greater retrieval evidence on retrieve compared to encode trials late in the stimulus interval. **(D)** We do not find a three-way interaction between instruction, distance, and time (*p* = 0.893). Error bars denote SEM. *∼* p *<* 0.1, ** p *<* 0.01, *** p *<* 0.001.

We next sought to investigate the effect of temporal overlap on retrieval. If greater spatiotemporal contextual overlap between two events promotes retrieval, we would expect to find greater evidence for a retrieval state on *near* compared to *far* trials. Moreover, to the extent that this retrieval occurs automatically, we would expect to find greater evidence for a retrieval state early in the stimulus interval. Although temporal distance could interact with instruction – evidence for a retrieval state may be particularly strong for *near* retrieve trials – given that temporal distance did not enhance memory for *near* List 1 objects on retrieve trials or impact memory for List 2 objects, we do not anticipate an interaction between temporal distance and instruction.

To investigate the effect of temporal distance on retrieval state evidence over time, we trained classifiers to discriminate encode vs. retrieve trials using the average z-power from four 500 ms time intervals across the 2000 ms stimulus interval. We conducted a repeated measures ANOVA in which retrieval evidence was the dependent variable and with factors of instruction (encode, retrieve), distance (*near, far*) and time interval (four 500 ms time intervals). We find a significant two-way interaction between distance and time interval (*F*_3,108_ = 5.585, *p* = 0.0013) whereby retrieval evidence is greater for *near* compared to *far* trials during the first two 500 ms time intervals (*near* vs. *far* : 0-500, *t*_36_ = 1.9781, *p* = 0.0556; 500-1000, *t*_36_ = 3.2961, *p* = 0.0022) and greater for *far* compared to *near* trials during the 1000-1500 ms time interval (*t*_36_ = -2.778, *p* = 0.0086). Retrieval evidence does not differ in the final 1500-2000 ms time interval (*t*_36_ = 0.3606, *p* = 0.7205). We also find a trending main effect of distance (*F*_1,36_ = 3.27, *p* = 0.0789; Figure 3B), with greater retrieval evidence for *near* compared to *far* trials. We find a significant two-way interaction between instruction and time interval (*F*_3,108_ = 5.125, *p* = 0.0024), whereby the largest differences in retrieval evidence between retrieve and encode trials occurs during the last two 500 ms time intervals (encode vs. retrieve: 0-500, *t*_36_ = -1.3956, *p* = 0.1714; 500-1000, *t*_36_ = -1.6667, *p* = 0.1043; 1000-1500, *t*_36_ = -4.255, *p* = 0.0001; 1500-2000, *t*_36_ = -4.3626, *p* =0.0001). We find a significant main effect of instruction (*F*_1,36_ = 21.31, *p <* 0.0001; Figure 3C), consistent with the results of the classifier trained on the full 2000 ms interval above. The twoway interaction between instruction and distance was not significant (*F*_1,36_ = 2.221, *p* = 0.1450) nor was the three-way interaction between instruction, distance, and time interval (*F*_3,108_ = 0.205, *p* = 0.8930; Figure 3D). Together, these results suggest that greater spatiotemporal contextual overlap induces automatic retrieval independent of the actual instruction to either encode or retrieve. Interestingly, we find greater retrieval evidence on *far* compared to *near* trials in the 1000-1500 ms interval. This effect is largely driven by the encode trials (Figure 3D) and is consistent with a suppression interpretation whereby automatic retrieval of the List 1 object is followed by top-down encoding of the List 2 object.

### Contribution of Serial Position to the Link Between Temporal Distance and Retrieval State

We find greater retrieval evidence for *near* compared to *far* trials, specifically early in the stimulus interval. However, due to the design of the experiment, *near* List 2 objects appear in early serial positions (primacy) and *far* List 2 objects appear in later serial positions (recency). Spectral signals are known to vary as a function of serial position (Sederberg et al., 2006; Serruya, Sederberg, & Kahana, 2014). Specifically, the neural serial position effect is characterized by increased high frequency (*>* 28 Hz) spectral power and decreased low frequency (*<* 28 Hz; Burke et al., 2014) spectral power for primacy relative to recency items. Thus, the dissociation in retrieval state evidence between *near* and *far* trials may be driven by a serial position effect rather than retrieval *per se*. To formally address this concern, we conducted a univariate analysis to assess serial position effects in both lists in our study, as well as a multivariate analysis to test the extent to which serial position effects contribute to the classifier results reported above, using List 1 as a control to assess serial position effects independent of both distance and instruction.

First, we tested for a neural serial position effect by assessing univariate differences in primacy (serial position 1-9) vs. recency (serial position 10-18) objects. We calculated the difference in spectral power between primacy and recency objects separately for List 1 and List 2 across 63 electrodes and 46 frequencies. We expected to find increases in power at high frequencies and decreases in power at low frequencies for primacy compared to recency objects in both lists. Indeed, our results mirror the classic serial position effect (Figure 4A). To test the serial position effects, we ran a repeated measures ANOVA on each list with frequency and electrode as factors with the dependent variable being the contrast of primacy vs. recency trials. To simplify the analysis, we collapsed the 46 frequencies into two bands, low frequency activity (LFA, *<* 28 Hz) and high frequency activity (HFA, *>* 28 Hz). For both List 1 and List 2 objects, we find a significant main effect of frequency (*F* ’s *>* 30, *p*’s *<* 0.0001), reflecting increased HFA and decreased LFA for primacy compared to recency objects. We also find an interaction between frequency and electrode in both lists (*F* ’s *>* 3.5, *p*’s *<* 0.0001) and a main effect of electrode in List 1 (*F*_62,2232_ = 4.753, *p <* 0.0001). These results reveal the classic serial position effect and suggest that these spectral differences may underlie the dissociation in retrieval state evidence between *near* and *far* List 2 trials.

**Figure 4.**
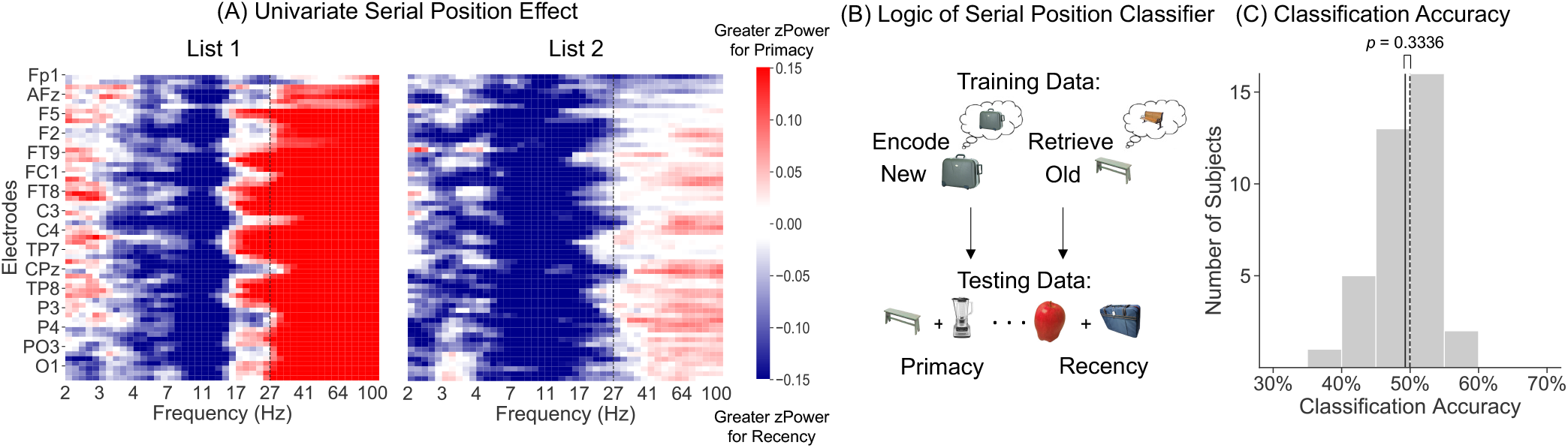
Serial Position Effects. **(A)** Electrode-frequency spectrogram of differences in spectral power between primacy and recency objects as a function of electrode (y-axis) and frequency (x-axis). Red indicates greater *z*-power for primacy objects (objects from serial positions 1-9), blue indicates greater *z*-power for recency objects (objects from serial positions 10-18). Spectrograms were generated for each subject and then averaged across subjects. Across both lists, we find increased high frequency activity (HFA) and decreased low frequency activity (LFA) for primacy compared to recency objects. The dashed line indicates the cutoff between HFA (*>* 28 Hz) and LFA (*<* 28 Hz). **(B)** To investigate whether the dissociation in retrieval state evidence between *near* and *far* trials is driven by serial position, we trained an L2-logistic regression classifier to discriminate encode vs. retrieve List 2 trials and tested the ability of this classifier to distinguish primacy from recency objects during List 1 trials. If serial position effects contribute to the dissociation between List 2 *near* and *far* objects, then the classifier should also be able to dissociate primacy and recency List 1 objects. **(C)** Mean classification accuracy across all subjects (solid vertical line) is shown along with a histogram of classification accuracies for individual subjects (gray bars) and mean classification accuracy for permuted data across all subjects (dashed vertical line). Mean classification accuracy for permuted data ranged from 49.70% to 50.12% across individual subjects (1000 permutations per subject). Mean classification accuracy was 49.29%, which did not differ significantly from chance (*p* = 0.3336).

To determine if the univariate serial position effects observed above contribute to the dissociation in retrieval state evidence between *near* and *far* trials, we conducted a multivariate pattern classification analysis in which we trained a classifier to discriminate encode vs. retrieve List 2 trials and tested this classifier to distinguish primacy from recency objects during List 1 (Figure 4B). The logic of this analysis is that if the classifier trained on the List 2 trials is dissociating *near* from *far* objects by virtue of their serial position (primacy vs. recency), this classifier should also be able to dissociate primacy and recency List 1 objects, given that List 1 and List 2 show comparable univariate serial position effects. That is, to the extent that the classifier is leveraging serial position effects to distinguish *near* from *far* List 2 objects, it should perform above chance (50%) when tested on List 1 primacy and recency objects.

We averaged spectral power across the stimulus interval (2000 ms) for both List 1 and List 2 trials. We trained a classifier to distinguish encode/retrieve List 2 and tested the classifier’s ability to discriminate primacy from recency List 1 trials. Mean classification accuracy was 49.29% (SD = 0.0437%), which did not differ significantly from chance (*t*_36_ = -0.98, *p* = 0.3336; Figure 4C). This suggests that the dissociation in retrieval evidence between *near* and *far* trials is unlikely to be driven by spectral differences across serial positions.

### Retrieval State Mechanisms

We have found an increase in retrieval state evidence when objects appear closer together in time. Although our hypothesis is that this dissociation reflects greater instantiation of a retrieval state, the classifier may be indexing retrieval success or retrieval effort as opposed to a general retrieval state or mode (Rugg & Wilding, 2000). Specifically, by virtue of the shorter temporal distance, retrieval success might be greater for *near* compared to *far* objects. Likewise, by virtue of the longer temporal distance, retrieval might be more effortful for *far* compared to *near* objects. In our previous classification analysis, the classifier was trained using data from both *near* and *far* trials, meaning that the dissociation between encode/retrieve trials, and consequently, *near* /*far* trials, could be based on information exclusively from either *near* or *far* trials. Put another way, the classifier may have learned to distinguish either encode and ‘retrieval success’ (i.e. *near* retrieve) trials or encode and ‘retrieval effort’ (i.e. *far* retrieve) trials. Therefore, to demonstrate that a general retrieval state or mode underlies the dissociation between *near* and *far* trials, we trained two separate classifiers to distinguish encode/retrieve using only *near* or only *far* trials, and tested the classifiers on the other held-out distance (*far* or *near*) trials. The logic is that to the extent that the dissociation between encode/retrieve is supported by the same mechanism on both *near* and *far* trials, classifiers trained on one distance should generalize – reflected by above chance (50%) performance – to the other distance. To the extent that the dissociation between encode/retrieve is driven either by retrieval success or retrieval effort, the classifiers should fail to generalize to the other distance.

We conducted a multivariate pattern classification analysis in which we trained a classifier on only *near* or *far* trials to discriminate encode vs. retrieve trials. We averaged spectral power across the stimulus interval (2000 ms) and used leave-one-run-out cross-validated classification. First, we trained a classifier to distinguish encode vs. retrieve List 2 *near* trials and tested the classifier on the List 2 *far* trials (Figure 5A). Mean classification accuracy was 52.62% (SD = 0.0529%), which was significantly greater than chance performance (*t*_36_ = 2.9734, *p* = 0.0052; Figure 5A). Next, we trained a classifier to distinguish encode vs. retrieve List 2 *far* trials and tested the classifier on List 2 *near* trials (Figure 5B). Mean classification accuracy was 53.33% (SD = 0.0588%), which was significantly above chance (*t*_36_ = 3.401, *p* = 0.0017; Figure 5B). Together, these results suggest that the dissociation in retrieval evidence between *near* and *far* trials is likely due to a retrieval state or mode rather than retrieval effort or retrieval success.

**Figure 5.**
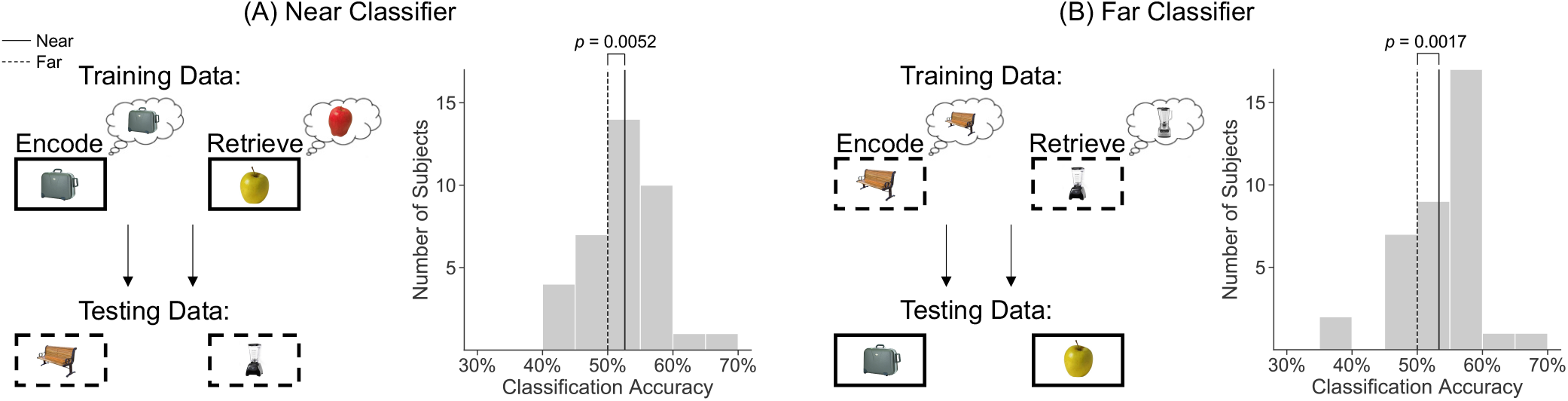
Cross Distance Mnemonic State Decoding. We trained two L2-logistic regression classifiers to discriminate encode vs. retrieve based on average spectral power for the 2000 ms stimulus interval with 63 electrodes and 46 frequencies used as features. For each classifier we show mean classification accuracy across all subjects (solid vertical line) along with a histogram of classification accuracies for individual subjects (gray bars) and mean classification accuracy for permuted data across all subjects (dashed vertical line). **(A)** We trained the classifier on only List 2 *near* trials and tested the classifier on List 2 *far* trials. Mean classification accuracy for permuted data ranged from 49.96% to 50.04% across individual subjects (1000 permutations per subject). Mean classification accuracy was 52.62%, which was significantly greater than chance performance (*p* = 0.0052). **(B)** We trained the classifier on only List 2 *far* trials and tested the classifier on List 2 *near* trials. Mean classification accuracy for permuted data ranged from 49.97% to 50.04% across individual subjects (1000 permutations per subject). Mean classification accuracy was 53.33%, which was significantly greater than chance performance (*p* = 0.0017).

## Discussion

Here we show that spatiotemporal contextual overlap between events selectively increases interference and induces automatic retrieval. We used scalp EEG to measure memory brain states in a task during which subjects were explicitly instructed to either encode the present event or retrieve a past, overlapping event. We find that temporal overlap selectively leads to interference for past events when the top-down goal is to encode the present event. We find that temporal overlap induces automatic retrieval independent from top-down demands to encode or retrieve. Critically, our results suggest that the retrieval state we observe is likely the result of a general retrieval ‘mode’ (Rugg & Wilding, 2000), rather than a reflection of retrieval success or effort. Collectively, these findings demonstrate a link between spatiotemporal context, interference, and memory brain states.

We find that greater temporal overlap between events leads to a selective memory deficit for a past event when the top-down demand is to encode the present event. Overlap between events can lead to both proactive interference, in which learning about a past event impairs memory for the present, and retroactive interference, in which learning about a present event impairs memory for the past (Underwood, 1948; Crowder, 1976). Here we find that greater temporal overlap between two events leads to an increase in retroactive interference; however, this increase is selective for conditions in which subjects’ top-down goal is to encode the currently presented stimulus. This result has striking similarity with retrieval induced forgetting (Anderson et al., 1994; Anderson & Spellman, 1995). In paradigms that produce retrieval induced forgetting, subjects retrieve a target (e.g. strawberry) based on a word stem (e.g. s___) and a cue (e.g. food) that is associated with other non-targets (e.g. tomato). Researchers theorize that cue driven retrieval of the non-target leads to suppression or inhibition which impairs subsequent memory for the non-target (*c*.*f*. Perfect et al., 2004). As the strength, typically framed in terms of semantic overlap, between non-target and cue increases, there is an increase in memory impairment, putatively due to stronger inhibition (Anderson et al., 1994). We extend these findings by showing that temporal overlap can likewise impair memory for non-targets, suggesting that greater temporal overlap may lead to inhibition of automatically retrieved items that are not goal relevant.

Although in our study we find that temporal overlap is detrimental to later memory, there is evidence that temporal overlap between events can facilitate behavior. When making inference judgments based on overlapping associations, participants perform better when the associations are studied closer in time (Zeithamova & Preston, 2017). In free recall, events presented close together in time are often remembered together (temporally clustered, Kahana, 1996; Long & Kahana, 2015) and overall memory performance increases as more events are temporally clustered (Sederberg, Miller, Howard, & Kahana, 2010; Healey, Crutchley, & Kahana, 2014). It is possible that in our study the explicit instruction to encode interrupts or prevents integration (Schlichting & Preston, 2015; Richter, Chanales, & Kuhl, 2016) between the two events, leading to worse memory for the past event. More generally, temporal overlap may have a differential impact on behavior depending on the type of memory assessment, as greater neural pattern similarity across events facilitates memory in free recall, but impairs memory for paired associates (El-Kalliny et al., 2019).

We find stronger induction of a retrieval state – specifically early in the stimulus interval and independent of top-down demands – when objects are closer together in time. We anticipated that greater temporal overlap would lead to increased retrieval on the basis of retrieved context theory. According to retrieved context theory (Howard & Kahana, 2002; Sederberg et al., 2008; Polyn et al., 2009; Lohnas & Kahana, 2014), spatiotemporal context is bound to items during study and used as a retrieval cue during test (Long et al., 2017), enabling items with overlapping spatiotemporal contexts to cue retrieval of one another (Manning et al., 2011). Consistent with retrieved context theory, we find more retrieval state evidence for objects with greater temporal overlap (*near* compared to *far* objects). Furthermore, this dissociation in retrieval state evidence is largest within the first 1000 ms following stimulus onset, suggesting that this retrieval is automatic. Automatic retrieval is thought to be a fast, bottom-up process that can occur without top-down control (Moscovitch, 1994). Our observation of elevated retrieval state evidence on *near* trials even when the instruction is to encode the present (or, conversely, when the instruction is to *not* retrieve the past), coupled with the early timing of this effect, strongly suggests that we are observing automatic retrieval driven by a bottom-up property of the object (e.g. its spatiotemporal contextual overlap with a past object) rather than a result of top-down demands. Interestingly, later in the stimulus interval we find a ‘flip’ in retrieval state evidence, whereby there is stronger retrieval state evidence for *far* compared to *near* trials. Although this pattern is evident when subjects are instructed to encode as well as to retrieve, the effect is relatively larger on encode trials. These results suggest that a present event may be encoded more strongly if its overlapping past event occurred recently in time; however, we did not find evidence that temporal overlap influences memory for the present event. Collectively, these findings indicate that memory brain states can rapidly change depending on both bottom-up and top-down influences.

The retrieval state driven by temporal overlap that we observe likely reflects a general retrieval ‘mode’ rather than serial position effects, retrieval success, or retrieval effort. We replicate the neural serial position effect whereby high frequency activity increases and low frequency activity decreases during primacy compared to recency items (Sederberg et al., 2006; Serruya et al., 2014); however, it is unlikely that these neural signals underlie the dissociation in retrieval state evidence between *near* and *far* trials. More critically, the dissociation between *near* and *far* trials could be the result of other retrieval processes rather than a more general retrieval ‘mode’ (Tulving, 1985; Rugg & Wilding, 2000). *Retrieval* as it stands is a broad concept and can encompass multiple different ‘sub-processes.’ We consider a retrieval state or mode as a content-independent process. Although typically retrieval mode has been considered within the framework of goal-directed or intentional remembering, we expect that a retrieval state can also be engaged automatically based on bottom-up inputs (as demonstrated in the current study) and may align or be synonymous with the internal axis of attention (Chun, Golomb, & Turk-Browne, 2011). This retrieval mode or state is thought to be distinct from retrieval ‘orientation’ in which specific cues or features are used to guide memory (Herron & Wilding, 2004; Hornberger, Rugg, & Henson, 2006a, 2006b). Finally, both retrieval state and orientation are separate from retrieval success and retrieval effort. After directing attention internally and orienting to particular cues to guide retrieval, an individual will either bring to mind the target item (retrieval success) or fail to bring to mind the target item, leading to effortful retrieval. The retrieval process that we observe in the current study likely reflects a general retrieval state given that a pattern classifier can distinguish memory encoding and retrieval across both *near* and *far* trials. If the processes underlying *near* and *far* trials were entirely related to retrieval success and retrieval effort, respectively, the pattern classifier would be unable to distinguish encoding and retrieval across these trials. This is not to say that there are not potential differences in terms of retrieval success or retrieval effort between *near* and *far* trials, only that there exist shared mechanisms which enables the pattern classifier to generalize across these trials. These results present an exciting avenue for future work to use multivariate approaches to further dissociate these different retrieval sub-processes and to more generally relate memory retrieval to internal attention.

Our results add to a growing body of work demonstrating the presence of neurally dissociable mnemonic states (Hasselmo et al., 2002; Hasselmo, 2005). Mnemonic states predict subsequent memory (Long & Kuhl, 2019), impact the cortical location of stimulus representations (Long & Kuhl, 2021), and can influence behavior and decision making (Duncan, Sadanand, & Davachi, 2012; Duncan & Shohamy, 2016; Patil & Duncan, 2018). Here we show that mnemonic states are influenced by bottom-up stimulus properties, or the features of a stimulus, in addition to explicit top-down demands. We expect that mnemonic states fluctuate based on both stimuli and goals – to the extent that events overlap, there is the potential for automatic retrieval and a shift into a retrieval state. Tracking mnemonic state fluctuations will be critical for understanding both how these states are induced and how these states in turn impact behavior.

In summary, we show that temporal overlap between events induces automatic retrieval and promotes interference. These findings are consistent with theoretical models which propose that temporal information can cue retrieval (Howard & Kahana, 2002) and behavioral findings that retrieving non-goal relevant information can lead to memory impairments (Anderson et al., 1994). More broadly, these findings point to a role for bottom-up stimulus features in driving mnemonic brain states.

## Acknowledgments

We thank Yuju Hong for assistance with data collection. Nicole Long is an iTHRIV Scholar. The iTHRIV Scholars Program is supported in part by the National Center for Advancing Translational Sciences of the National Institutes of Health under Award Numbers UL1TR003015 and KL2TR003016.

